# Modeling osteoporosis to design and optimize pharmacologic therapies comprising multiple drug types

**DOI:** 10.1101/2021.12.17.473190

**Authors:** David J. Jörg, Doris H. Fürtinger, Alhaji Cherif, David A. Bushinsky, Ariella Mermelstein, Jochen G. Raimann, Peter Kotanko

**Affiliations:** Biomedical Modeling and Simulation Group, Global Research and Development, Fresenius Medical Care Germany, Bad Homburg, Germany; Renal Research Institute, New York, New York, United States of America; Department of Medicine, University of Rochester School of Medicine and Dentistry, Rochester, New York, United States of America; Icahn School of Medicine at Mount Sinai, New York, New York, United States of America

## Abstract

For the treatment of postmenopausal osteoporosis, several drug classes with different mechanisms of action are available. Since only a limited set of dosing regimens and drug combinations can be tested in clinical trials, it is currently unclear whether common medication strategies achieve optimal bone mineral density gains or are outperformed by alternative dosing schemes and combination therapies that have not been explored so far. Here we develop a mathematical framework of drug interventions for postmenopausal osteoporosis that unifies fundamental mechanisms of bone remodeling and the mechanisms of action of four drug classes: bisphosphonates, parathyroid hormone (PTH) analogs, sclerostin inhibitors and receptor activator of NF-*κ*B ligand (RANKL) inhibitors. Using data from several clinical trials, we calibrate and validate the model, demonstrating its predictive capacity for complex medication scenarios including sequential and parallel drug combinations. Via simulations, we reveal that there is a large potential to improve gains in bone mineral density by exploiting synergistic interactions between different drug classes, without increasing the total amount of drug administered.

## INTRODUCTION

Osteoporosis, a disease characterized by porous bone prone to fractures, affects hundreds of millions of people worldwide [1, 2]. Most recent estimates place the global annual incidence of bone fragility fractures at 9 million in the year 2000 [1]; projections for the year 2050 suggest between 7 and 21 million annual hip fractures [3]. Osteoporosis-associated bone fractures lead to disabilities, pain and increased mortality [1]. In the United States, medical cost for osteoporosis including inpatient, outpatient and long-term care costs have been estimated at US$17 billion in 2005 [4]; in the European Union, the total cost of osteoporosis including pharmacological interventions and loss of quality-adjusted life years (QALYs) is projected to rise from about EUR 100 billion in 2010 to EUR 120 billion in 2025 [5].

Osteoporotic bone is the consequence of an imbalance of continuous bone resorption and bone formation, which—under close to homeostatic conditions—has the function to remove microfractures and renew the structural integrity of bone. Postmenopausal women are particularly at risk of osteoporosis: The rapid decline of systemic estrogen levels after menopause and other aging-related effects such as increased oxidative stress contribute to or drive the development of osteoporosis [6, 7]. Moreover, osteoporosis can be a sequela of diseases affecting bone metabolism and remodeling such as primary hyperparathyroidism or gastrointestinal diseases [8]. Osteoporosis can also be a side effect of treatments for other diseases; as a prime example, glucocorticoid administration is the most common cause of secondary osteoporosis [9]. Over the last decades, an array of different osteoporosis treatments have emerged, from simple dietary supplementations such as calcium and vitamin D to specialized drugs targeting bone forming and resorbing cells and related signaling pathways [10]. This entails a plethora of different medication options including a large number of possible dosing schemes and combinations of drugs, administered in sequence or in parallel. Due to the huge number of such treatment schemes and the required time from study inception to completion, very few of them have been clinically tested so far when compared to the total number of available options.

Concomitant with the development of new osteoporosis drugs, mathematical and biophysical modeling approaches capturing bone-related physiology have advanced our quantitative understanding of the biological principles governing bone mineral metabolism, bone turnover and development of osteoporosis. Pioneering work by Lemaire *et al*. [11] describes the dynamics of bone-forming and resorbing cell populations coupled through signaling pathways and could qualitatively reproduce the effects of senescence, glucocorticoid excess and estrogen and vitamin D deficiency on bone turnover. Since then, compartment-based descriptions of the mineral metabolism, bone forming and resorbing cell populations and related signaling factors have elucidated the role of essential regulatory mechanisms underlying mineral balance and bone turnover [11–21]. Coarse-grained as well as detailed spatially extended descriptions of bone geometry have also addressed the effects of mechanical forces and the propagation of the multicellular units responsible for bone turnover [22–26] as well as the influence of secondary diseases such as multiple myeloma [27]. Detailed models of bone remodeling and calcium homeostasis have become versatile and widely used tools in hypothesis testing, such as the seminal model by Peterson and Riggs [15], which includes submodels for various organs such as gut, kidney and the parathyroid gland. Pharmacokinetic and pharmacodynamic (PK/PD) models of therapeutic interventions have mostly focused on capturing the mechanisms of action of a single or a few drugs and testing their dosing regimens [28–35]. Recent modeling efforts including multiple drugs have also started addressing the effects of combination therapies on bone-forming and resorbing cells [36], pointing out the need for corresponding model frameworks to include clinically relevant variables like bone mineral density (BMD) and bone turnover biomarkers (BTMs). An integrated mathematical framework for multiple drugs, which can also be used to quantitatively predict the effects of drug combinations in sequence and in parallel is not yet available.

Building on established mechanisms of bone turnover, we here present a quantitative model of bone turnover and postmenopausal osteoporosis treatment, unifying the description of multiple classes of drugs with different mechanisms of action, namely bisphosphonates, parathyroid hormone (PTH) analogs, sclerostin antibodies and receptor activator of NF-*κ*B ligand (RANKL) antibodies. We calibrate the model using published population-level data from several clinical trials and assess its ability to predict the outcome of previously conducted clinical studies based on the medication scheme alone. We then use the model to demonstrate how medication schemes involving drug combinations can be optimized for a given medication load and discuss future model extensions.

## MECHANISMS OF BONE TURNOVER AND ITS REGULATION

Our model is based on a small set of key principles of bone turnover, which we briefly recapitulate here (Fig. 1). As a composite tissue comprising hydroxyapatite, collagen, other proteins and water [37], bone is constantly turned over to renew its integrity and remove microdamage, at an average rate of about 4% per year in cortical bone and about 30% per year in trabecular bone [38].

**Figure 1.**
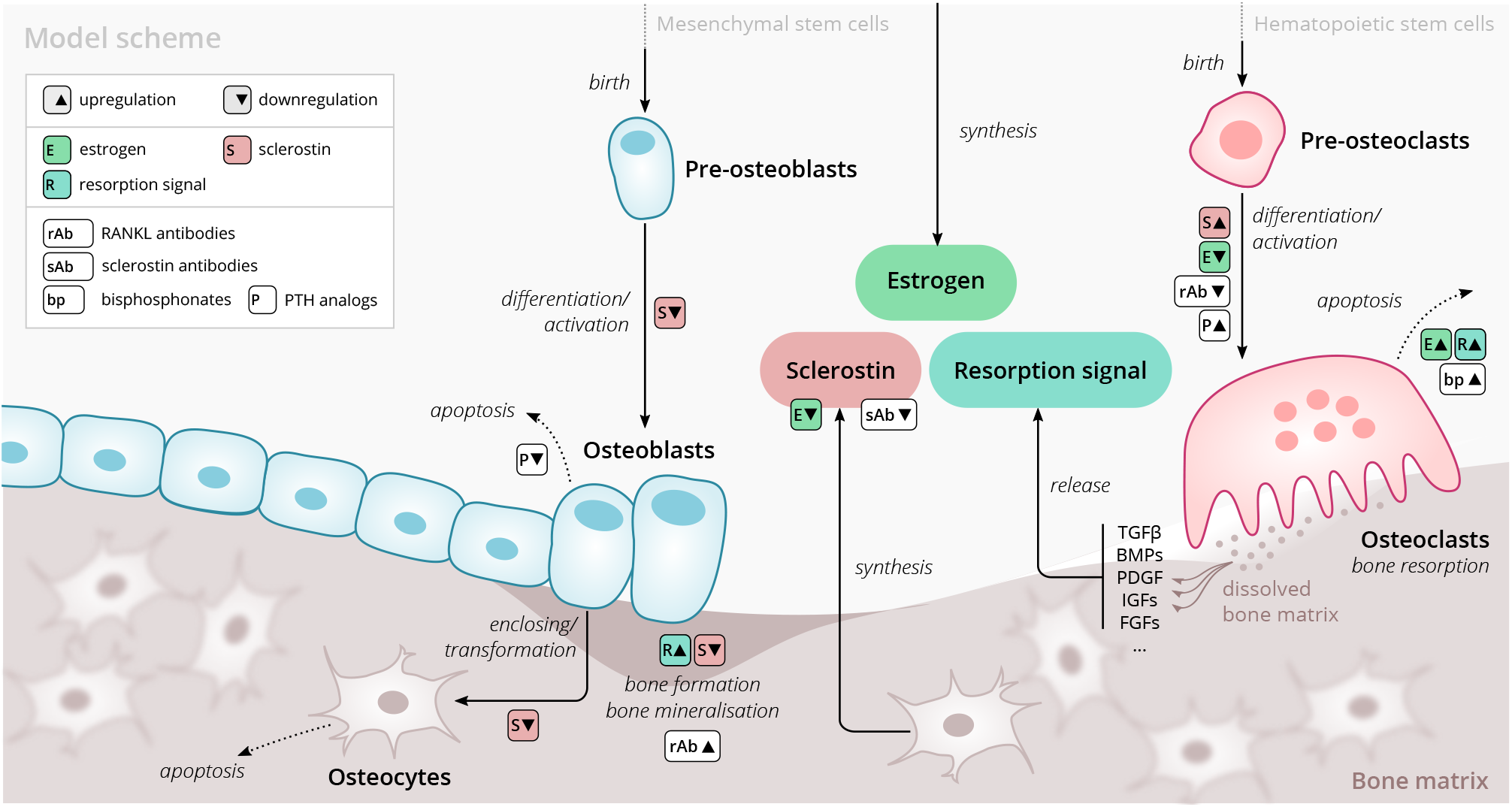
Schematic of the osteoporosis model describing the cell dynamics and signaling pathways within a ‘representative bone remodeling unit (BRU)’. Regulatory interactions between different model components are indicated by colored boxes (see legend). Abbreviations: TGF*β*—transforming growth factor *β*, BMP—bone morphogenetic protein, PDGF—platelet-derived growth factor, IGF—insulin-like growth factor, FGF—fibroblast growth factor.

### Bone-resorbing and forming cells

Bone resorption is performed by osteoclasts, multinucleated cells formed through the differentiation and fusion of their immediate precursors (pre-osteoclasts), which are derived from pluripotent hematopoietic stem cells via the myeloid lineage [39]. Osteoclasts attach to bone tissue and resorb it through the secretion of hydrogen ions and bone-degrading enzymes [40], which leads to the release of minerals and signaling factors stored in the bone matrix. New bone is formed by osteoblasts, a cell type derived from mesenchymal stem cells via several intermediate states that give rise to preosteoblasts and finally osteoblasts [41]. Groups of osteoblasts organize into cell clusters (osteons) and collectively lay down an organic matrix (osteoid) which subsequently becomes mineralized over the course of months. Osteoblasts that are enclosed in the newly secreted bone matrix become osteocytes, non-dividing cells with an average lifespan of up to several decades. Osteoclasts and osteoblasts organize into spatially defined local clusters termed ‘bone remodeling units’ (BRUs) (Fig. 1), in which osteoblasts replenish the bone matrix previously resorbed by osteoclasts with a delay of several weeks. In cortical bone, the outer protective bone layer, BRUs migrate as a whole in ‘tunnels’ whereas within the inner cancellous bone, BRUs propagate on the surfaces of the trabeculae, renewing the bone matrix in the process [41].

### Signaling pathways

The differentiation and activity of osteoclasts and osteoblasts are regulated through several signaling pathways and hormones; recent reviews provide comprehensive descriptions of the various pathways [42]. Osteoclast formation and activity are prominently regulated by receptor activator of NF-*κ*B ligand (RANKL) and macrophage colony-stimulating factor (M-CSF) synthesized by bone marrow stromal cells. RANKL binds to receptor activator of NF-*κ*B (RANK) on osteoclast precursors and promotes their differentiation into mature osteoclasts; osteoprotegerin (OPG) acts as a decoy receptor for RANKL and thus inhibits bone resorption [39, 43]. When laying down new bone, osteoblasts store signaling factors in the bone matrix, including transforming growth factor beta (TGF*β*), bone morphogenetic protein (BMP), insulin growth factors (IGFs), platelet-derived growth factor (PDGF) and fibroblast growth factors (FGFs) [44]. Upon bone resorption, these factors are released and regulate cell fates and activity of osteoblasts and osteoclasts, thereby coupling bone resorption and formation [41, 45]. Osteocytes secrete sclerostin, a Wnt inhibitor interfering with extracellular binding of Wnt ligands [46]. Sclerostin inhibits bone formation and promotes resorption via downregulation of osteoblastogenesis and upregulation of osteoclastogenesis [47, 48]. Since bone also acts as a mineral reservoir for the body, regulators of calcium homeostasis such as parathyroid hormone (PTH) and vitamin D also strongly affect the balance of bone formation and resorption alongside the intestinal absorption and renal reabsorption of calcium [49].

### Estrogen

The sex hormone estrogen inhibits bone resorption by inducing apoptosis of osteoclasts [50] and lowering circulating sclerostin levels [51]. The rapid decline of estrogen levels after menopause is one known cause of postmenopausal osteoporosis [6].

## RESULTS

### Model overview

The primary purpose of our model is to provide an efficient representation of bone turnover on multiple time scales from weeks to decades that allows for the quantitative description of drug interventions. Of particular interest are the consequences of pharmacologic therapies on long-term dynamics of the bone mineral density (BMD) in specific bone types and biochemical markers of bone formation and resorption. To this end, we considered a minimal set of physiologically relevant dynamic components (Fig. 1) that are sufficient to capture a large range of clinically observed population-level data on drug interventions. Thus, our model describes a ‘representative bone remodeling unit (BRU)’ that abstracts from the vast set of intricate regulatory mechanisms underlying calcium homeostasis or the complex bone geometry.

Our model comprises the following dynamic components to describe the bone turnover through a representative BRU: cell densities of (i) pre-osteoclasts, (ii) osteoclasts, (iii) pre-osteoblasts, (iv) osteoblasts, (v) osteocytes; (vi) sclerostin concentration; (vii) total bone density and (viii) bone mineral content. The BMD is given by the product of bone density and bone mineral content. Osteoblasts and osteoclasts can undergo apoptosis and are derived from pre-osteoblasts and pre-osteoclasts, respectively, with differentiation rates that depend on regulatory factors such as estrogen and sclerostin (Fig. 1). Pre-osteoblasts and pre-osteoclasts are formed at constant rates and undergo apoptosis. These progenitor populations provide a dynamic reservoir for rapid differentiation and activation of osteoblasts and osteoclasts, respectively, which can be temporarily depleted if stimulated by a drug intervention. Osteocytes are derived from osteoblasts and provide a source of sclerostin, which has a regulatory effect on osteoblasts, osteoclasts and thus, bone density change. The gain and loss rates of bone density are proportional to the density of osteoblasts and osteoclasts, respectively. The bone mineral content has a steady state whose level can be temporarily shifted through drug administration, effectively accounting for more complex underlying dynamics such as promotion of secondary mineralization. All rates of cell formation, differentiation, apoptosis and bone formation and resorption generally depend on the concentration of sclerostin, estrogen and a ‘resorption signal’. These dependencies also implicitly account for regulation of bone remodeling via other routes, e.g., the RANK– RANKL–OPG pathway. The effects of aging and the onset of menopause are represented through an age-dependent serum estrogen concentration, which has been determined from the literature [52] (Appendix A). The resorption signal corresponds to the melange of signaling factors stored in the bone matrix. Therefore, its release is proportional to the rate of bone resorption. The serum concentration of bone turnover markers (BTMs) such as the resorption marker C-terminal telopeptide (CTX), the formation markers procollagen type 1 amino-terminal propeptide (P1NP) and bone-specific alkaline phosphatase (BSAP) were identified with elementary functions of the bone resorption and formation rates in the model (Appendix A).

We extended this core model of long-term bone turnover by a dynamic description of the mechanisms of action of several drug classes used in osteoporosis treatment: RANKL antibodies (denosumab), sclerostin antibodies (romosozumab), bisphosphonates (alendronate and others) and PTH analogs (teriparatide) (Appendix B). We also included blosozumab, another sclerostin inhibitor, which was investigated in osteoporosis trials but not approved for osteoporosis treatment at the time the present work was conducted. PTH is known to exert anabolic or catabolic effects depending on whether administration is intermittent or continuous [53, 54]; PTH description in our model is restricted to the anabolic administration regimes relevant for osteoporosis treatment. A schematic overview of all model components, mechanisms and regulatory interactions is given in Fig. 1; a detailed formal description of the model and its extensions is provided in Appendices A and B.

### Capturing clinical study results with the model

The model and the corresponding medication modules rely on an array of physiological parameters (rates of cell formation, differentiation and death, concentration thresholds for signaling activity, medication efficacies and half-lives etc.) many of which are not directly measureable. However, clinical measurements on physiological responses to medications with different mechanisms of action provide a wealth of indirect information about time scales of bone turnover and regulatory feedbacks. We calibrated the model using published clinical data from various seminal studies on both (i) longterm BMD age dependence and (ii) the response of BMD and BTMs to the administration of different drugs (see Table 1 for a comprehensive list of data sources). Although BMD constitutes the major target variable of our model, the dynamics of BTM concentrations carry important complementary information about the mode of action of the administered drugs (antiresorptive, anabolic and combinations) that crucially informs the calibration procedure. To allow the model to capture the effects of medications as physiologically sensible modulations of the age-dependent bone mineral metabolism, we created hybrid datasets each of which comprised both aging-related BMD changes and the response to a treatment (Materials and Methods and Supplementary Fig. 1D).

**Table 1.**
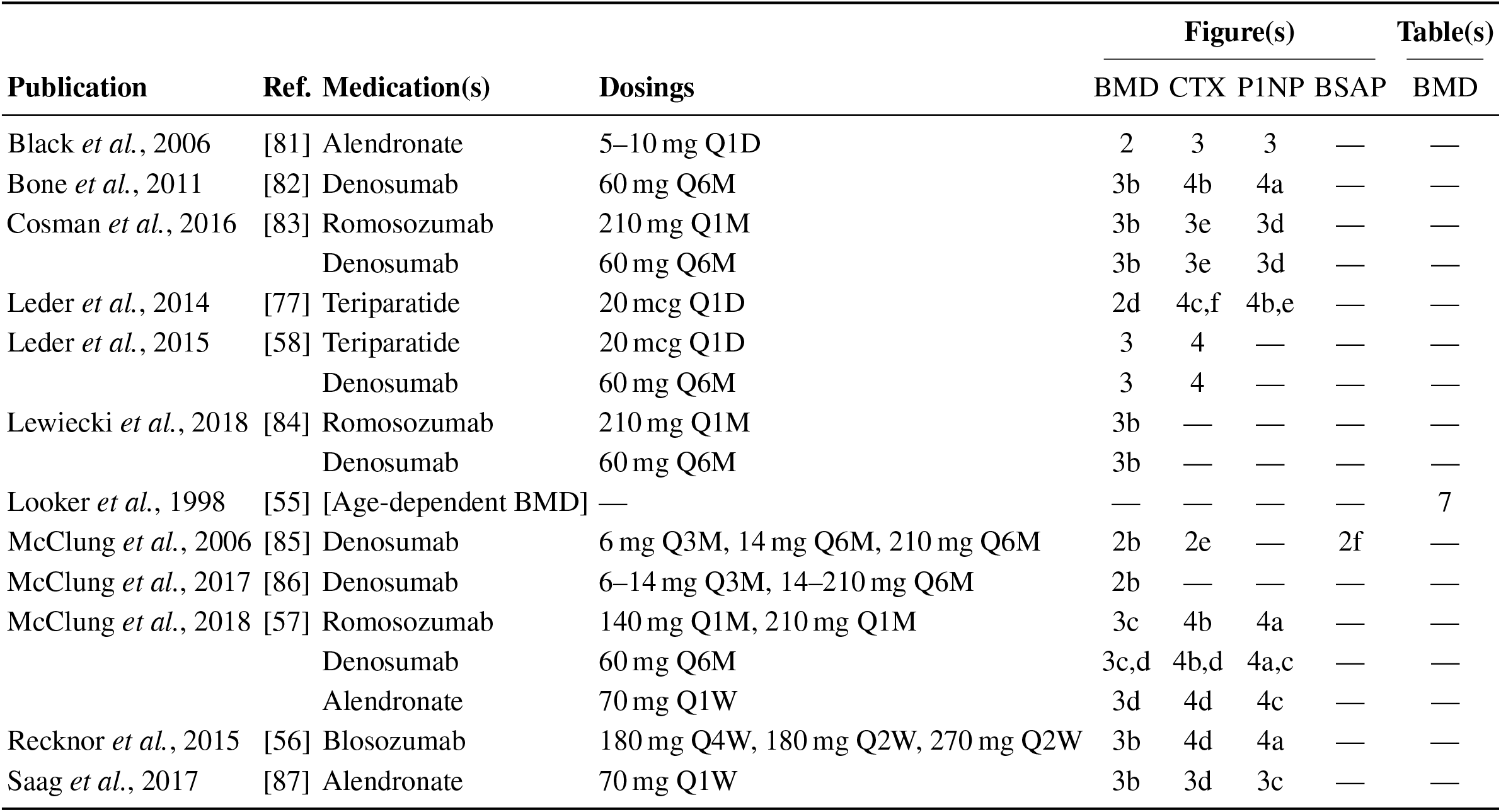
Data sources used to calibrate and validate the model. Columns titled ‘Figure(s)’ indicate the plot panels in the respective publication that were digitized. BMD always refers to total hip bone mineral density.

We then determined a single set of model parameters through a simultaneous fit of the free 31 model parameters to capture a specified set of hybrid aging/treatment datasets containing different drug responses (Appendix C). Without constraining the average rate of skeletal bone turnover, model calibration yielded an inferred value of about 6% per year on average, of the same order of values reported for cortical bone, which constitutes about 75% of the skeleton [38]. The model was able to capture the BMD and BTM dynamics across all calibration datasets with remarkable accuracy (Supplementary Fig. 2), despite the model’s structural simplicity. To quantify the goodness of the fit, we computed the mean absolute percentage error (MAPE) between model simulations and clinical data; the MAPE for BMD was consistently below 1% for all calibration datasets (Table 2), indicating an excellent agreement between model and data. The qualitative behavior of BTMs (i.e., the direction of their excursions from baseline) was captured correctly in all calibration datasets, indicating an adequate description of the drugs’ mode of action in the model; relative deviations in the total magnitude of BTM excursions observed for some datasets were mostly due to slight offsets in the timing of peaks and troughs and low absolute values of the respective BTM concentrations, as highlighted by comparing different goodness measures (Appendix C and Table 2).

**Table 2.** Goodness of fit measures for calibration and validation datasets. Mean absolute percentage error (MAPE) and Windowed minimal absolute percentage error (WMAPE) as defined in Eqs. (C3) and (C4), respectively. The column ‘Shown in’ indicates the main text or supplementary figure in the present paper that shows the respective simulation and data plot.

| Medication(s) | Data ref. | MAPE |  |  |  | WMAPE |  |  |  | Shown in |
| --- | --- | --- | --- | --- | --- | --- | --- | --- | --- | --- |
|  |  | BMD | CTX | P1NP | BSAP | BMD | CTX | P1NP | BSAP |  |
| Calibration datasets |  |  |  |  |  |  |  |  |  |  |
| Alendronate 5-10mg Q1D | [81] | 0.9% | 7.0% | 21.1% | — | 0.6% | 4.7% | 13.5% | — | SFig. 2 |
| Alendronate 70mg Q1W | [87] | 0.5% | 34.4% | 13.7% | — | 0.2% | 20.3% | 10.2% | — | SFig. 2 |
| Blosozumab 180mg Q4W | [56] | 0.7% | 16.5% | 23.8% | — | 0.4% | 10.8% | 18.7% | — | SFig. 2 |
| Blosozumab 270mg Q2W | [56] | 0.3% | 26.8% | 13.2% | — | 0.2% | 15.8% | 4.8% | — | SFig. 2 |
| Denosumab 14mg Q3M → Denosumab 60mg Q6M | [86] | 0.3% | — | — | — | 0.1% | — | — | — | SFig. 2 |
| Denosumab 14mg Q6M | [85] | 0.2% | 73.9% | — | 17.0% | 0.0% | 46.3% | — | 0.2% | SFig. 2 |
| Placebo | [56] | 0.4% | 7.3% | 15.4% | — | 0.3% | 7.2% | 15.3% | — | SFig. 2 |
| Teriparatide 20mcg Q1D | [77] | 0.4% | 17.1% | 8.5% | — | 0.2% | 9.1% | 1.6% | — | SFig. 2 |
| Teriparatide 20mcg Q1D → Denosumab 60mg Q6M | [58] | 0.4% | 65.5% | — | — | 0.2% | 8.6% | — | — | SFig. 2 |
| Validation datasets |  |  |  |  |  |  |  |  |  |  |
| Alendronate 70mg Q1W → Romosozumab 140mg Q1M → Denosumab 60mg Q6M | [57] | 0.5% | 25.4% | 20.0% | — | 0.3% | 17.1% | 3.6% | — | Fig. 3 |
| Blosozumab 180mg Q2W | [56] | 0.3% | 21.3% | 20.3% | — | 0.1% | 13.2% | 12.6% | — | Fig. 3 |
| Denosumab 60mg Q6M → Teriparatide 20mcg Q1D | [58] | 0.4% | 103.1% | — | — | 0.3% | 44.6% | — | — | Fig. 3 |
| Placebo | [57] | 0.6% | 6.2% | 10.5% | — | 0.5% | 6.2% | 10.5% | — | Fig. 3 |
| Placebo → Denosumab 60mg Q6M | [57] | 0.6% | 13.2% | 10.1% | — | 0.4% | 5.1% | 7.4% | — | Fig. 3 |
| Teriparatide 20mcg Q1D + Denosumab 60mg Q6M → Denosumab 60mg Q6M | [58] | 0.7% | 183.7% | — | — | 0.4% | 66.4% | — | — | Fig. 3 |
| Alendronate 70mg Q1W → Romosozumab 140mg Q1M → Placebo | [57] | 0.5% | 22.9% | 16.8% | — | 0.4% | 15.1% | 4.2% | — | SFig. 3 |
| Placebo → Denosumab 60mg Q6M | [83] | 0.5% | 81.9% | 9.6% | — | 0.0% | 44.4% | 4.0% | — | SFig. 3 |
| Placebo → Denosumab 60mg Q6M | [84] | 0.4% | — | — | — | 0.1% | — | — | — | SFig. 3 |
| Placebo → Denosumab 60mg Q6M | [86] | 0.6% | — | — | — | 0.3% | — | — | — | SFig. 3 |
| Romosozumab 210mg Q1M → Alendronate 70mg Q1W | [87] | 1.0% | 30.0% | 37.1% | — | 0.5% | 19.4% | 11.7% | — | SFig. 3 |
| Romosozumab 210mg Q1M → Denosumab 60mg Q6M | [83] | 1.2% | 92.5% | 22.4% | — | 0.6% | 50.7% | 6.9% | — | SFig. 3 |
| Romosozumab 210mg Q1M → Denosumab 60mg Q6M | [84] | 1.0% | — | — | — | 0.5% | — | — | — | SFig. 3 |
| Romosozumab 210mg Q1M → Denosumab 60mg Q6M | [57] | 0.6% | 14.2% | 57.8% | — | 0.3% | 6.5% | 15.9% | — | SFig. 3 |
| Romosozumab 210mg Q1M → Placebo | [57] | 0.7% | 15.0% | 53.3% | — | 0.5% | 10.2% | 18.5% | — | SFig. 3 |

After obtaining the reference parameter set, we sought to validate the calibrated model by assessing its ability to predict the effects of drug dosing schemes that had not been used for calibration. Model validation included complex sequential and parallel drug combinations and therefore challenged the model to predict the effects of treatment schemes beyond those used in calibration (Table 1). To this end, the model received only drug dosing information used in the respective clinical trials but was not informed by BMD or BTM measurements, which instead it had to predict. With the single set of previously determined parameters, the model showed a remarkable capacity to quantitatively forecast the effects of a multitude of medication schemes, both during treatment and follow-up periods (Fig. 2 and Supplementary Fig. 3). Even in scenarios including sequential treatments with up to three different drug types and parallel treatments with two different drugs, respectively, the model was able to predict the complex progression of both bone mineral density and biomarker levels with a high degree of accuracy (Fig. 2). Across all validation datasets, MAPEs for BMD were consistently below 1.5% (Table 2), indicating an excellent predictive capacity of the model. In summary, this validation provided a strong corroboration of the model’s capacity to capture the physiological dynamics of bone turnover and the mechanisms of action of various drugs relevant to osteoporosis treatment, using a single set of model parameters.

**Figure 2.**
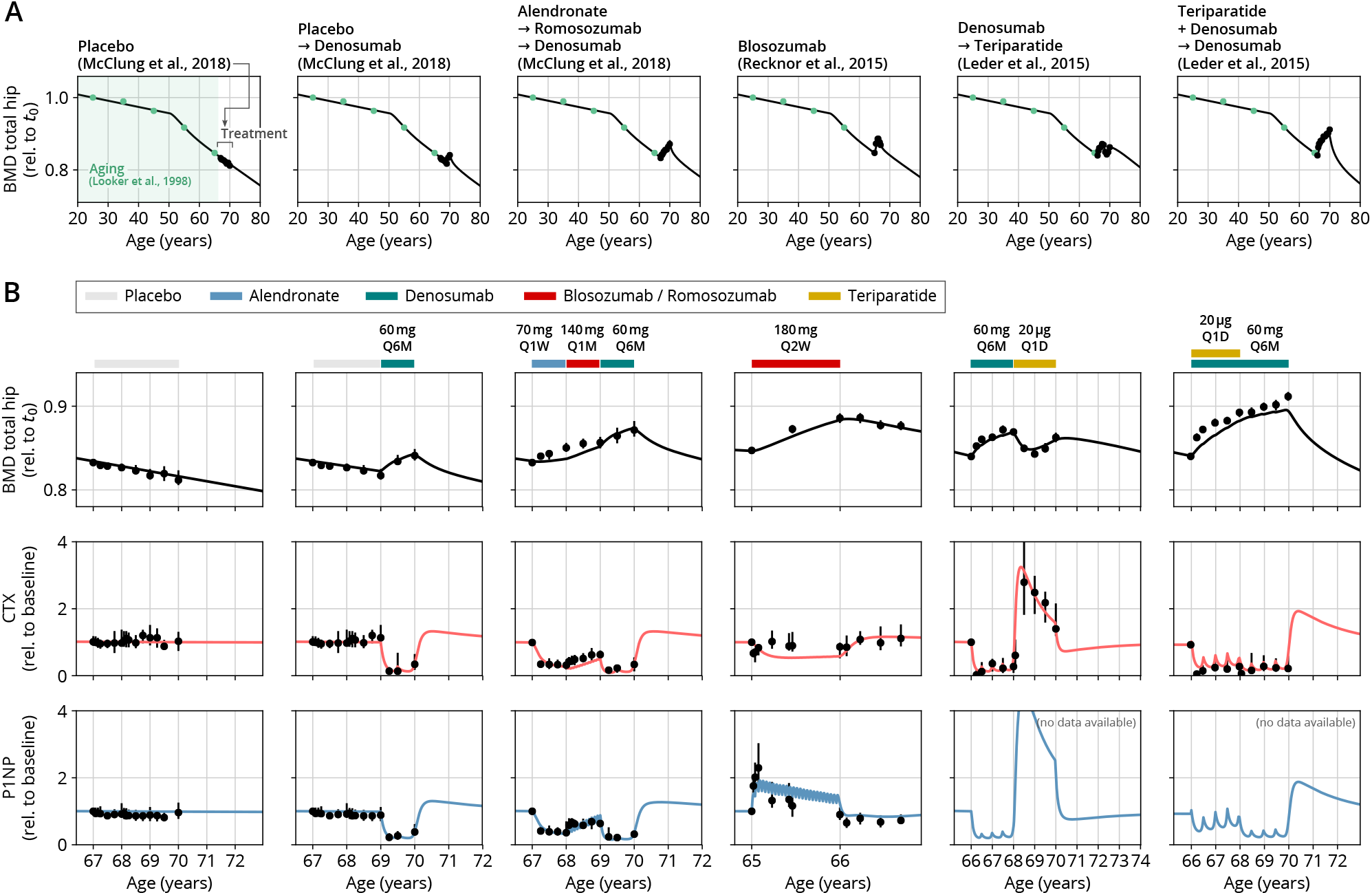
With a single set of parameters, the calibrated model can quantitatively predict the effects of various drugs in different dosing regimens, alone and in combination. (A) Comparison of simulated total hip bone mineral density (BMD, black curves) and clinical data (dots) including aging behavior (green dots) and treatment behavior (black dots) of various sequential drug treatments including denosumab, romosozumab, alendronate and teriparatide. Hybrid aging/treatment datasets were created combining data from Looker *et al*., 1998 [55] (aging dataset, green dots in panel A; in total *N* = 3251 subjects 20 years and older) as well as Recknor *et al*., 2015 [56] (blosozumab 180mg Q2W: *N* = 25), McClung *et al*., 2018 [57] (placebo/deno.: *N* = 18, alendro./romo./deno.: *N* = 21), and Leder *et al*., 2015 [58] (deno./teri.: *N* = 27, teri.+deno./deno.: *N* = 23) (treatment datasets, black dots in panels A and B) as indicated, see Materials and Methods. (B) Zoom into the treatment regions shown in panel A including BMD (black) and baseline changes of the bone resorption marker C-terminal telopeptide (CTX, red) and the bone formation marker procollagen type 1 amino-terminal propeptide (P1NP, blue). Colored bars above the plots indicate the medication scheme (see legend). Data points show population averages; average types and error bar types as reported in the respective original publication. In both panels, BMD is displayed as a fraction of its value at *t*_0_ = 25 years. Figure continued in Supplementary Fig. 3.

### Testing alternative treatment schemes

Having established the predictive capacity of the model for the considered medications, we aimed to utilize the model to study and optimize hypothetic drug dosing regimens. As an example, we considered a sequential treatment with three drugs of different types: the bisphosphonate alendronate, the sclerostin inhibitor romosozumab and the RANKL inhibitor denosumab. In a clinical trial reported by McClung *et al*. [57], the sequence alendronate (70 mg per week for one year), followed by romosozumab (140 mg per month for one year), followed by denosumab (60 mg per 6 months for one year) had been studied (Fig. 2). However, in principle there are six different sequences in which these drugs can be administered: ARD, ADR, DAR, DRA, RAD and RDA (A: alendronate, R: romosozumab, D: denosumab). A priori, it is not obvious whether synergistic or antagonistic interactions between these drugs and the physiological state in which they leave the patient may lead to a differential short and long-term evolution of BMD and biomarkers between different medication sequences. Probing all six sequences in a clinical trial would present a time- and resource-consuming endeavour and inevitably expose part of the study population to suboptimal treatment schemes. Instead, we probed these different treatment options using the present model (Fig. 3A). To assess the predicted clinical success of different sequences, we compared two clinically relevant outcomes across different schemes: (i) the maximum achieved BMD increase (as compared to baseline at treatment start) irrespective of when it occurred (Fig. 3B) and (ii) the residual long-term effects of treatment on BMD as monitored by the relative BMD ten years after treatment end (Fig. 3C).

**Figure 3.**
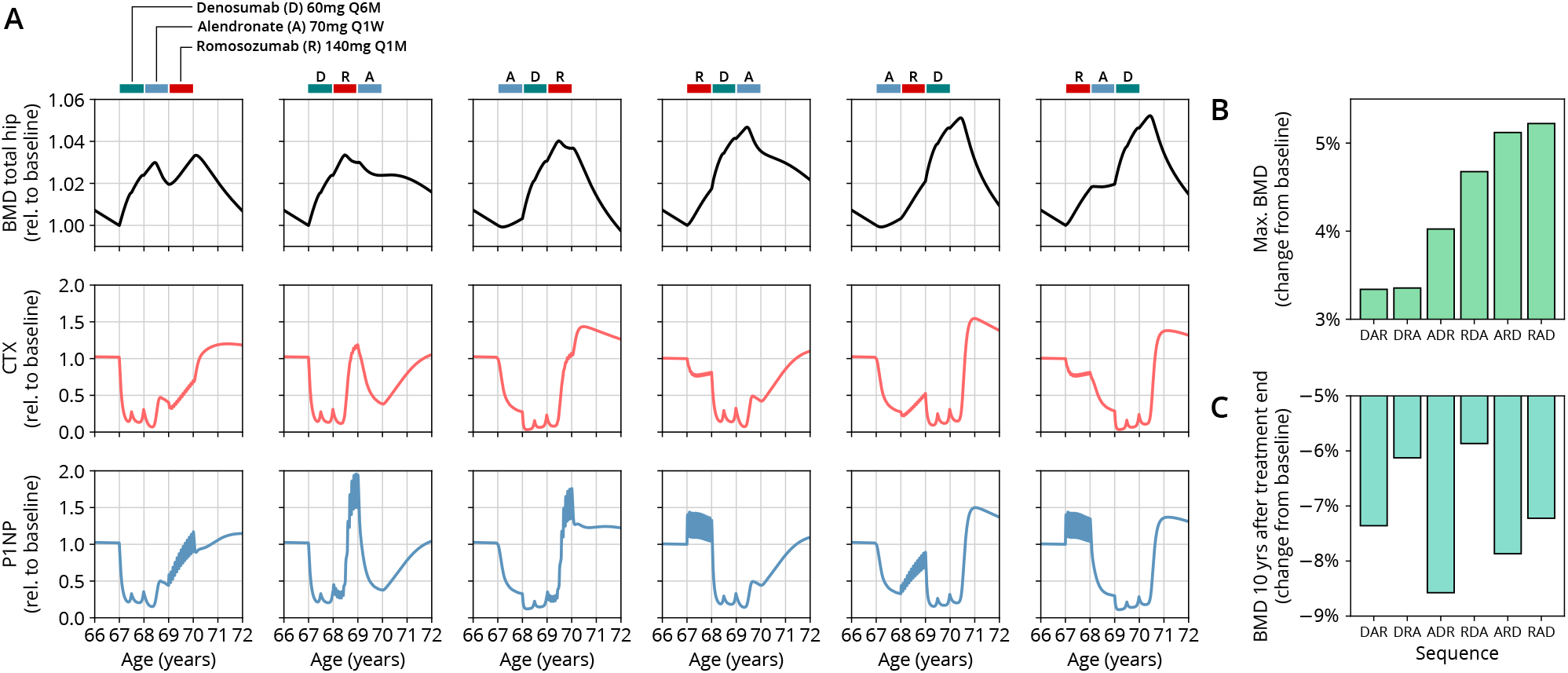
The model predicts differential outcomes for different sequences of the same drugs at constant total medication load. (A) Simulated progression of BMD and CTX and P1NP concentrations for different sequences (columns) of the three drugs denosumab (D), alendronate (A) and romosozumab (R) as indicated. Simulated treatment starts at age 67. The total amount of drug administered is identical among columns. Clinical results on the sequence ARD (column 5) were reported in McClung *et al*., 2018 [57], see also Fig. 2. (B) Maximum simulated BMD (relative to baseline at treatment start) achieved during the course of treatment for different drug sequences. (C) Simulated BMD ten years after treatment end (relative to baseline at treatment start) for different drug sequences.

Indeed, we found that the outcomes of different medication sequences were markedly different despite the same total amount of drug administered (Fig. 3A). Some sequences (such as ARD and RAD) reached a considerably higher maximum BMD during the course of the simulated treatment, which allowed us to rank treatments according to maximum BMD gain (Fig. 3B). Notably, while some sequences were superior to others as measured by the maximum BMD increase during treatment, they performed markedly worse (as compared to, e.g., DRA and RDA) with regard to long-term BMD evolution as predicted by model simulations (Fig. 3C). This behavior suggests that short-term BMD gains may be limited as a proxy for the clinical benefit of a treatment as a whole. Within our modeling scheme, the explanation for this behavior is found in differing ‘rebound’ effects after treatment end: simulated drug-mediated inhibition of osteoclastogenesis leads to a build-up of an undifferentiated osteoclast precursor pool. After treatment end, this precursor pool becomes licensed to differentiate and rapidly gives rise to a large active osteoclast population, leading to accelerated resorption of the bone matrix that had been built up during treatment. In this paradigm, specific drug sequences lead to an attenuation of this effect, e.g., by enhancing osteoclast apoptosis during such a ‘rebound’ phase, thereby modulating bone turnover in the long run.

In summary, our model analysis suggests considerable potential in the improvement of dosing regimens and drug sequencing in osteoporosis treatment, especially combination therapies, to achieve an optimal effect for a given medication load. These improvements are possible because the mechanisms of action of one drug may act either favorably or adversely on the state of the bone mineral metabolism left behind by the preceding treatment with another drug.

## Discussion

We have introduced a biomedical modeling framework that can quantitatively capture and predict the progression of a multitude of drug-based osteoporosis treatments in postmenopausal women. Our model is built on a small set of essential mechanisms of bone turnover; the effectivity of this approach suggests that—despite the complexity of the bone mineral metabolism—the dynamics relevant for osteoporosis medications can be condensed into only a few components describing (i) the cell fates and activity responses of osteoblasts, osteoclasts and osteocytes, (ii) temporarily exhaustible ‘reservoirs’ for rapid bone turnover activity (embodied through the precursor cell populations in the present model) and (iii) a few essential regulatory feedbacks through hormones and signaling factors such as estrogen, sclerostin and bone-matrix derived factors.

The parsimonious nature of the model allowed us to determine a single parameter set with which the model is able to capture the bone mineral density and bone turnover markers of a multitude of clinical studies. Notably, using this single set of parameters, the model can also predict the effects of a broad range of dosing regimens and complex drug combination therapies beyond those used for calibration. This corroborates the model’s predictive capacity which may be used to inform the design of future clinical trials. Nonetheless, despite the simplified nature of the model, the dimensionality of the parameter space is considerable. Mesoscopic parameters (like concentration thresholds for signaling factors) were inferred through the calibration procedure from BMD and BTM dynamics alone and can, in principle, only be pinpointed by independent measurements. Hence, used as a predictive tool, general quantitative limitations of the model have to be considered, especially when extrapolating into extreme dosing regimes, dosing frequencies or age regions beyond the validated ones.

Exemplary model predictions suggest a large potential for the development of optimized combination therapies involving different drug types and the optimization of existing treatment schemes. These may range from a simple rearrangement of a sequence of drugs at given total drug doses (as shown in this paper) to complex interwoven or cyclic administration schemes that exploit synergistic effects between different medication types. Model simulations extrapolating the long-term development after treatment end suggest that medication schemes eliciting a rapid BMD increase are not necessarily accompanied by a sustained elevation of the BMD. Instead, some initially successful treatment schemes may lead to a ‘rebound’ effect of accelerated bone loss after treatment end, a prediction that cautions against using short-term BMD increases as the sole proxy for treatment success. Such extrapolations into regimes inaccessible to clinical studies highlight the potential role of the model in taking into account long-term treatment success when optimizing treatment schemes.

Although emphasis has been placed on postmenopausal osteoporosis, the most widespread type of osteoporosis, the generic manner in which the model represents bone remodeling and the effects of medications renders it a general platform for the study of treatments that can be adapted to other types of primary and secondary osteoporosis. The modular nature of the model enables future extensions; besides additional medication types, these may include the effects of comorbidities that elicit osteoporosis or interact with it (such as primary and secondary hyperparathyroidism), treatments of comorbidities that contribute to osteoporosis (such as glucocorticoid therapy), related lifestyle-dependent factors such as physical activity, smoking and alcohol consumption, the effects of dietary supplementation of osteoporosis treatment through calcium and vitamin D and effects of microgravity on bone, as experienced by astronauts on extended missions in space. Clinical data used to calibrate such additional extensions typically carry information about bone turnover mechanisms already present in the model; therefore model extensions also present a chance to further pinpoint model parameters and to revalidate the extended model framework. Thus, our model can serve as a quantitative starting point for the forecast of drug-based therapies of osteoporosis but also highlights the role of mechanistic mathematical descriptions in understanding the biological principles underlying drug action.

## MATERIALS AND METHODS

### Hybrid aging/treatment datasets

To create hybrid aging/treatment datasets, we merged a dataset comprising the BMD age dependence from Looker *et al*. [55] with different clinical study datasets containing the BMD response to various medications (Table 1). The aging dataset from Looker *et al*. [55] consisted of mean total femur BMD measurements in non-Hispanic white, non-Hispanic black and Mexican American women, reported in ten-year age bins ranging from 20 to 80 years and older. We used bin averages as proxy BMD indicators for the center of the respective age window (Supplementary Fig. 1B). Rescaling the reported means for the three ethnic groups to their value for the earliest age bin revealed that relative changes in BMD were remarkably consistent among ethnic groups (Supplementary Fig. 1C) despite differing absolute baselines. Therefore, and since the model only addresses relative BMD changes, we resorted to the dataset with the largest underlying study population for calibration, which was the dataset comprising the non-Hispanic white female study population. Datasets on the response to medications from clinical trials on romosozumab, blosozumab, denosumab, alendronate and teriparatide consisted of study population averages of total hip BMD and serum concentrations of one or more bone turnover markers (CTX, P1NP, BSAP) during the treatment and if available, during a follow-up period. Reported study population averages on the respective quantities were digitized directly from the data figures in the corresponding publications (Table 1).

To merge aging and treatment datasets, the BMD from treatment datasets was rescaled such that the BMD baseline at treatment start corresponds to the linearly interpolated age-dependent BMD at treatment start. The treatment start was placed at the average age of the study population upon study start (rounded to full years) as reported in the respective publication (Supplementary Fig. 1D). BTM measurements were normalized to baseline values.

## ACKNOWLEDGEMENTS

We thank Friederike E. Thomasius for critical comments on the manuscript.

## AUTHOR CONTRIBUTIONS

D.J.J., D.H.F. and P.K. designed the research; D.J.J. developed the model, performed data extraction and data analysis; D.H.F. and A.C. provided input for modeling; D.A.B. and P.K. provided biological and clinical expertise; A.M. and J.G.R. contributed to data extraction; D.J.J. wrote the first draft of the manuscript; all authors reviewed and edited the manuscript.

## Appendix A: Model of long-term bone remodeling and osteoporosis

The model of bone remodeling underlying the present description of osteoporosis and its treatment was built with the aim to provide a minimal set of dynamic components necessary to quantitatively capture population averages of both aging-related changes in bone turnover and the response to osteoporosis medications with different mechanisms of action. The model is partitioned into a core model describing the patient physiology and separate extensions for different drug classes.

The model is compartment-based and describes average cell densities, bone densities and average concentrations of signaling factors within a ‘representative bone remodeling unit (BRU)’, i.e., a fictitious BRU corresponding to an average over the considered bone type. All model variables are treated as non-dimensional quantities, i.e., the model only addresses relative changes in all variables, which simplifies the model structure and reduces the number of free parameters.

### Model description

The model describes the dynamics of the cell densities of pre-osteoclasts (*ρ*_C*_), pre-osteoblasts (*ρ*_B*_), osteoclasts (*ρ*_C_), osteoblasts (*ρ*_B_), osteocytes (*ρ*_Y_) as well as functional sclerostin levels (*s*), total bone density (*ρ*_b_) and bone mineral content (*c*_b_). Functional estrogen levels (*e*) are provided as an explicitly age-dependent function, described further below. Moreover, the model includes a ‘resorption signal’ (*r*) corresponding to the composite concentration of bone matrix-derived signaling factors (TGF*β*, BMPs, PDGF, IGFs and FGFs) released upon bone resorption. Sclerostin, estrogen and the resorption signal act as regulatory factors that modulate the rates of cell proliferation, differentiation and apoptosis as well as their bone forming and resorbing activity; for notational convenience, they are summarized in the vector ***ϕ*** = (*s, e, r*). Rates that depend on ***ϕ*** are denoted with a tilde.

The dynamics of the cell density variables is given by

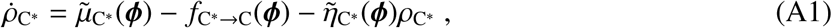

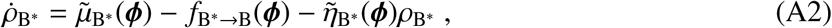

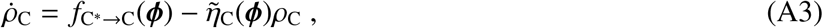

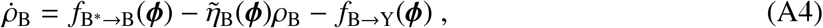

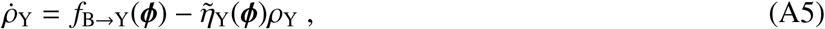

where dots denote time derivatives, 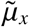 denotes the formation rate of cell population *x* and 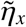 denotes its apoptosis rate. Differentiation or conversion from one cell type to another is described by the absolute differentiation rates *f*_*x*→*y*_ of cell population *x* giving rise to cells of type *y*; they have the generic form

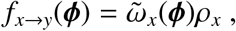

with 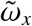 denoting the differentiation rate. All rates are functions of the regulatory factors ***ϕ*** as indicated.

Sclerostin is synthesized by osteocytes; the dynamics of Sclerostin levels is given by

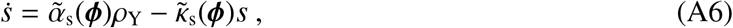

where 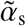 and 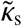 denote the synthesis and degradation rate, respectively.

The dynamics of the total bone density (BD) follows

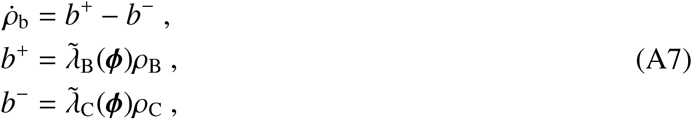

where *b*^+^ and *b*^−^ are the absolute rates of bone formation and resorption, respectively, with 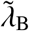 and 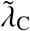 being the formation and resorption rates per unit osteoblast and osteoclast density, respectively.

The bone mineral content (BMC) follows the dynamics

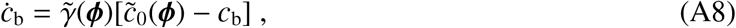

where 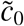 is the steady-state homeostatic bone mineral content and 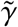 is the equilibration rate.

Based on the physiological foundations summarized in the Section ‘Mechanisms of bone turnover and its regulation’ in the main text, we now specify the functional form of all rates. Upregulation and downregulation through a regulatory factor with concentration *x* are described by a multiplicative or additive contribution *g*^+^(*x*/*x*_0_) or *g*^−^(*x*/*x*_0_), respectively, where *x*_0_ is the threshold concentration at which half-effect is reached (EC_50_) and where *g*^±^ are monotonic, saturating functions of the Hill-type [59],

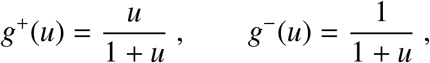

Using these conventions, the dependencies of rates on regulatory factors are given by

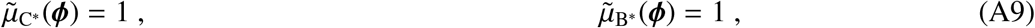

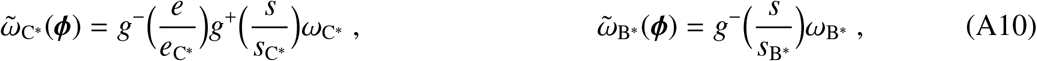

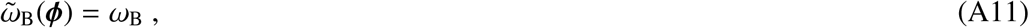

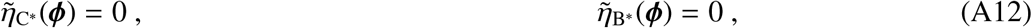

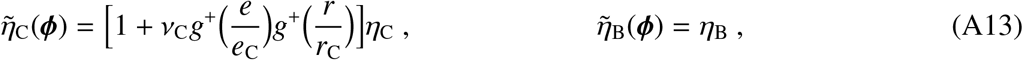

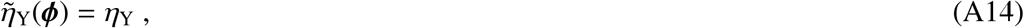

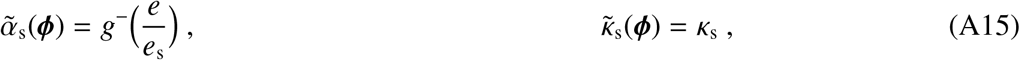

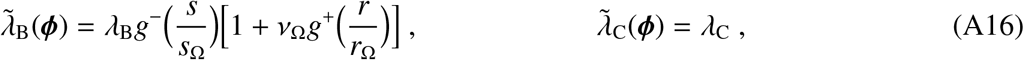

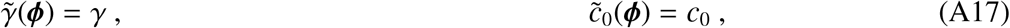

where rates without a tilde denote model parameters. A full list of parameters introduced here and their description is provided in Table 4. This regulatory scheme comprises a multitude of interactions, many of which are simplified effective representations of indirect molecular mechanisms (e.g., through the RANK–RANKL–OPG pathway as described below). In the model building process, the effects of additional regulatory elements (not presented here) were probed and found to be nonessential within the scope of the present modeling aim or non-identifiable with regard to related parameter values. Specific choices for regulatory interactions were partly based on insights from animal and culture studies and are motivated as follows.

Eqs. (A9): Pre-osteoclasts and pre-osteoblasts are produced at constant rates, a simplifying assumption reflecting the fact that the main function of these populations in the present context is to provide a dynamic reservoir for the rapid supply with active osteoclasts and osteoblasts, respectively.

Eqs. (A10): Estrogen suppresses pre-osteoclast to osteoclast differentiation; a consequence of suppression of RANKL expression [60]. Sclerostin upregulates pre-osteoclast to osteoclast differentiation; a consequence of upregulation of RANKL expression and downregulation of OPG expression [61]. Sclerostin downregulates osteoblastogenesis; a consequence of the inhibition of osteoblast differentiation mediated by bone morphogenetic protein 2 (BMP2) and Wnt3a and possibly other pathways [62, 63].

Eq. (A11): Sclerostin downregulates osteoblast to osteocyte conversion [64].

Eq. (A13): Estrogen and the resorption signal induce osteoclast apoptosis; estrogen has been reported to induce osteoclast apoptosis both directly and mediated by TGF*β* [50, 65]; TGF*β* has been shown to upregulate Bim, a member of the (pro-apoptotic) Bcl2 family [45].

Eqs. (A15): Estrogen reduces sclerostin production [51].

Eqs. (A16): Sclerostin downregulates bone formation [64, 66]. The resorption signal upregulates bone formation; a consequence of TGF*β*1 enhancing bone collagen synthesis [67]; furthermore, TGF*β*1, skeletal BMPs and IGF1 have been reported to inhibit collagenase-3 expression in osteoblasts [67–69].

Estrogen concentration is described through its explicit age dependence. Clinical data reported by Sowers *et al*. [52] were used to construct a function capturing the key features of age-dependent decline of serum estradiol levels,

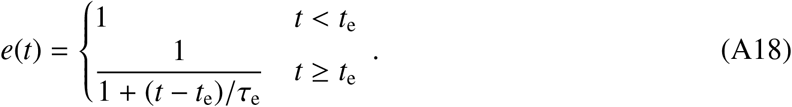

where *t* is time. The parameters *t_e_* and *τ*_e_, denote the age at the onset of estradiol decline and a characteristic time scale of the decline, respectively. The time scale *τ*_e_ was determined using a fit of the function to the data reported in Ref. [52] (Supplementary Fig. 1A and Table 4).

The resorption signal *r* corresponds to the concentration of bone matrix-derived signaling factors released upon bone resorption. Assuming a release rate proportional to the bone resorption rate *b*^−^ and first-order degradation, we consider a highly simplified dynamics of the type 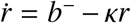, where *κ* is an effective average degradation rate of the components of the resorption signal. Given that the time scale of degradation, *κ*^−1^, is much shorter (minutes to hours) than the time scale of osteoclast formation and death (weeks), the instantaneous concentration can be approximated to always follow its steady state, *r* ≈ *b*^−^/*κ*, which is proportional to the osteoclast density, *r* ∝ *b*^−^ ∝ *ρ*^C^, via Eq. (A7). Since the resorption signal acts as a regulator of bone formation and is rescaled by individual concentration thresholds (see Eqs. A9–A17), the proportionality constant can be absorbed in these thresholds, which enables us to set *r* = *ρ*^C^. Thus, the resorption signal concentration is approximated by the osteoclast density, so that no additional dynamic variable is required.

### Description of bone mineral density, bone turnover rate and bone turnover markers (BTMs)

To compare the model output to clinical data, we relate model variables to clinical observables frequently measured in clinical trials such as bone mineral density and established biomarkers of bone turnover. The bone mineral density follows from the model state as the product of total bone density and bone mineral content,

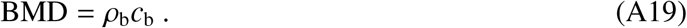

In our model, levels of bone turnover markers (BTMs) such as the bone formation markers procollagen type 1 amino-terminal propeptide (P1NP) and bone-specific alkaline phosphate (BSAP) and the bone resorption marker C-terminal telopeptide (CTX) are related to the rates *b*^+^ and *b*^−^ of bone formation and resorption, respectively, see Eqs. (A7). Here we relate the bone turnover markers P1NP, BSAP and CTX to bone turnover rates by power laws with marker-specific exponents,

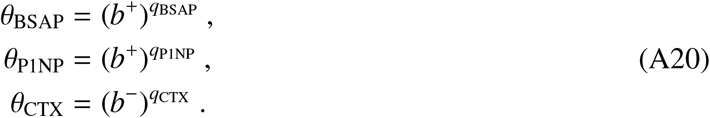

The exponents *q_x_* are obtained as fit parameters using clinical trial data, as described further below.

## Appendix B: Model extensions for medications

We include a dynamic description of the mechanism of action of several drug classes through separate model extensions, which depend on the functional drug concentration. While the pharmacodynamic description is drug-specific, the pharmacokinetics of all drugs described here follows a simple first-order kinetics with drug-individual half-lives. A patient’s systemic concentration of a medication is represented by an effective (dimensionless) variable *ψ* that indicates the relative concentration of the medication. Typically, *ψ* is given in multiples of a threshold that parametrizes the effect of the drug (such as EC_50_)—the precise interpretation of *ψ* depends on the model extension that describes the pharmacodynamics of the drug, see Section ‘Pharmacodynamics for specific medications’.

### Pharmacokinetics

The pharmacokinetics of a drug *x* that is administered in intervals of weeks or months is described by two parameters: the efficacy *E_x_* and the half-life *T_x_*. Given repeated administrations with doses *c*_1_, …, *c_n_* at times *t*_1_, …, *t_n_*, the efficacy-weighted concentration variable of the drug *x* therefore follows the exponential kinetics

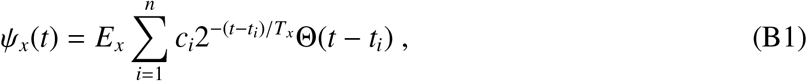

where Θ is the Heaviside function, defined by

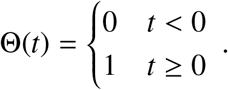

Drugs that are administered more frequently (e.g., daily or weekly) are more efficiently captured in a quasi-continuous scheme. The dynamics of BMD and BTM levels is much more inert than such fast administration/degradation dynamics and is well-described by their effective average action. In this quasi-continuous scheme, the drug is considered to be administered at a given average rate for a specified amount of time, so that its concentration evolves according to the dynamic equation

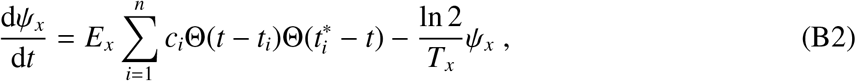

with initial condition *ψ_x_*(*t*)|_*t*→∞_ = 0, where *c_i_* are doses per unit time, and where *t_i_* and 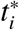 are the start and end times of a treatment period, respectively^1^.

Different drugs *x*_1_, *x*_2_, … of the same class (sclerostin inhibitors, bisphosphonates, PTH analogs etc.) are considered through an effective concentration equivalent that is the sum of the efficacy-weighted doses of different drugs,

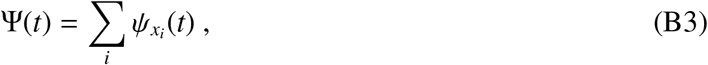

where the *ψ_xi_* are given by Eq. (B1) in the case of discrete doses and Eq. (B2) in the case of quasi-continuous dosing.

### Pharmacodynamics for specific medications

We now introduce separate model extensions that embody the essential mechanisms of action of different drug classes.

#### RANKL antibodies

Denosumab is a monoclonal antibody (mAb) that binds with affinity to receptor activator of NF-*κ*B ligand (RANKL) and blocks its interactions with RANK [70], which in turn, decreases osteoclast formation [39]. Moreover, denosumab has been suggested to enable increased bone tissue mineralization which leads not only to a halt of the BMD decline but also to a long-term increase in BMD [31]. We include these mechanisms in our model through a modification of the reference pre-osteoclast differentiation rate *ω*_C*_ to include a downregulation by denosumab and the steady-state BMC *c*_0_ to include an upregulation through denosumab in Eqs. (A10) and (A17), respectively:

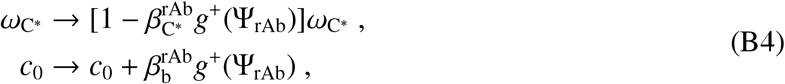

where Ψ_rAb_ is the RANKL antibody concentration in multiples of the half maximal effective concentration (EC_50_), determined through Eqs. (B1) and (B3). For simplicity, EC_50_ for the regulation of both differentiation and mineralization are taken to be identical, which is justified *a posteriori* by showing its effectivity in approximating the drug action. The scaling factors 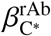 and 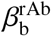 parametrize the respective maximum effect strength and are subject to the constraints 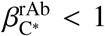 and 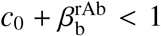 to ensure positive rates and BMCs between 0 and 100%.

#### Sclerostin antibodies

Romosozumab and blosozumab are monoclonal antibodies that bind to sclerostin and prevent its inhibitory effects on bone formation [56, 57, 71]. (Note that blosozumab was not approved for osteoporosis treatment at the time this manuscript was written.) Accordingly, we represent the mechanism of action of sclerostin antibodies by adding a new variable *s*\* corresponding to the level of antibody-bound sclerostin and adding the dynamics of antibody-binding and unbinding in Eq. (A6),

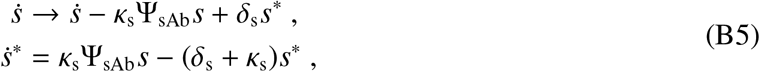

where Ψ_sAb_ is the effective sclerostin antibody concentration equivalent determined through Eqs. (B1) and (B3). Here, *κ*_s_ denotes the sclerostin degradation rate and *δ_s_* is the sclerostin/antibody binding rate; this parametrization implies that Ψ_sAb_ is given in multiples of the effective antibody levels needed to achieve a binding rate equal to the unperturbed degradation grade of sclerostin.

#### Bisphosphonates

Bisphosphonates (like alendronate, ibandronate, risedronate and zoledronate) bind to hydroxyapatite on the bone surface, thereby preventing osteoclasts from bone resorption; they further inhibit osteoclast-mediated bone resorption by promoting osteoclast apoptosis [72, 73]. To simplify the model extension, we effectively represent the mechanism of action of bisphosphonates through the upregulation of the osteoclast apoptosis rate in Eq. (A13),

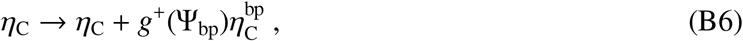

where Ψ_bp_ is the effective bisphosphonate concentration equivalent determined through Eqs. (B2) and (B3) and 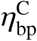 is the maximum additional apoptosis rate caused by the presence of bisphosphonates.

#### PTH analogs

Teriparatide and abaloparatide are recombinant human parathyroid hormone (PTH) analogs [74, 75]. PTH is known to have an anabolic effect on bone if administered intermittently while exerting a catabolic effect if administered continuously [53, 54]. Here we consider an effective representation of the action of PTH in the anabolic regime only; this leads to a highly simplified and efficient description of the effective action of PTH therapies on bone turnover. However, it implies that the scope of our model is restricted to anabolic administration schemes and cannot be expected to yield correct results if probed in inappropriate regimes. In the anabolic regime, teriparatide down-regulates osteoblast apoptosis [76]. Moreover, bone turnover markers show a marked increase early after treatment start but decline while drug administration remains unaltered [77], see Supplementary Fig. 2; such an effect is achieved in our model by rapid upregulation of osteoclast/osteoblast differentiation. We therefore also include a regulatory effect on osteoclast differentiation (which indirectly affects osteoblast differentation as well),

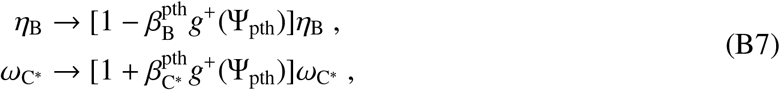

where Ψ_pth_ is the effective teriparatide concentration equivalent determined through Eqs. (B2) and (B3) and the parameters 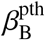 and 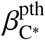 parameterize the maximum effect strength of osteoblast apoptosis and pre-osteoclast differentiation, respectively.

## Appendix C: Simulations and parameter fits

### Simulation protocol for aging and treatment

A model simulation was implemented in Python using standard NumPy and SciPy packages [78, 79] and solved using SciPy’s ‘solve_ivp’ function with BDF solver. To compare how the model predicts the bone turnover dynamics of a hybrid aging/treatment dataset (see Materials and Methods and Supplementary Fig. 1D) for a given set of model parameters, simulations were structured as follows. Drug dosing information of the corresponding dataset was provided to the model through the set of administered doses and the administration times: For the case of discrete dosing, dosing information consisted of doses *c_i_* and administration times *t_i_* entering Eq. (B1) (used for the drugs blosozumab, romosozumab and denosumab). For the case of quasi-continuous dosing, dosing information consisted of doses per unit time *c_i_* and time windows 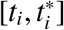 entering Eq. (B2) (used for the drugs alendronate and teriparatide).

The model was initialized at *t* = 0 in its steady state for all dynamic variables (i.e., the state for which all time derivatives are zero); except for the bone density *ρ*_b_, which does not possess a unique steady state and which was set to unity to represent peak bone density. We then simulated the model until well after the treatment period. All aging-related effects in the model were mediated by explicitly time-dependent auxiliary functions, as explained in the Section ‘Dynamic description of the model’. To compare simulation results with clinical data, all relevant model variables were rescaled such that the first recorded data point in the corresponding clinical data set coincided with the corresponding time point in the simulation, so that relative changes from a reference time point could be compared.

### Parameter fits

To systematically fit model parameters, we defined a cost function that takes into account multiple fit quantities depending on their availability in the datasets. For a hybrid dataset *α* and clinical quantity *β* (BMD and serum levels of CTX, P1NP and BSAP), we defined the distance function

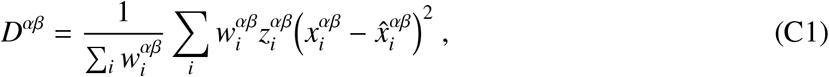

where *x_i_* denotes the clinical data point the time point *i*, 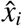 denotes the respective simulated data point, *w_i_* denotes the relative weight of the respective data point depending on its certainty and *z_i_* denotes the relative weight depending on the time interval represented by the respective data point. For BTMs, we used the weights 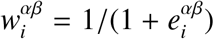 where 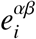 is the mean of the upper and lower error bar of the respective quantity *β*; for the BMD, we used unit weights 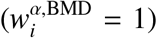. To account for the fact that time intervals between datapoints may vastly differ (e.g., between the coarse sampled aging dataset and the densely sampled treatment datasets), we included an interval-dependent weighting factor *z_i_*, such that each data point was weighted by the average distance to its neighboring data points: The time interval-related weighting factors 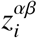 were defined as *z_i_* = (*δ*_*i*–1_ + *δ_i_*)/2 (1 ≤ *i* ≤ *n*), where *δ_i_* = *t*_*i*+1_ – *t_i_* (1 ≤ *i* ≤ *n* – 1), *δ*_0_ = *δ*_1_, *δ_n_* = *δ*_*n*–1_ and *t_i_* denotes the time point of measurement *i*.

We defined the combined cost function over all considered datasets as

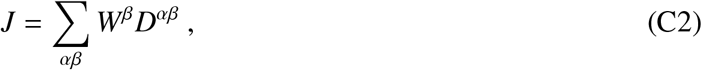

where *W^β^* is an additional weighting factor that determines the relative importance of the different fit quantities in the cost function (Table 3).

**Table 3.** Values of fit weights *W^β^* used in Eq. (C2).

| **Weight** | **Value** |
| --- | --- |
| $W^{(\text{BMD})}$ | 300 |
| $W^{(\text{CTX})}$ | 1 |
| $W^{(\text{PINP})}$ | 1 |
| $W^{(\text{BSAP})}$ | 1 |

In a first step, all fit parameters were manually adjusted for the model to exhibit a roughly sensible aging behavior. In a second step, a selected subset of hybrid aging/treatment datasets (see Materials and Methods and Supplementary Fig. 1D) were used to fit all free model parameters. To perform the fits, we used a Covariance Matrix Adaptation Evolution Strategy (CMA-ES) via the Python package ‘pycma’ [80]. The full list of fit parameters including their final fit values is given in Table 4.

**Table 4.**
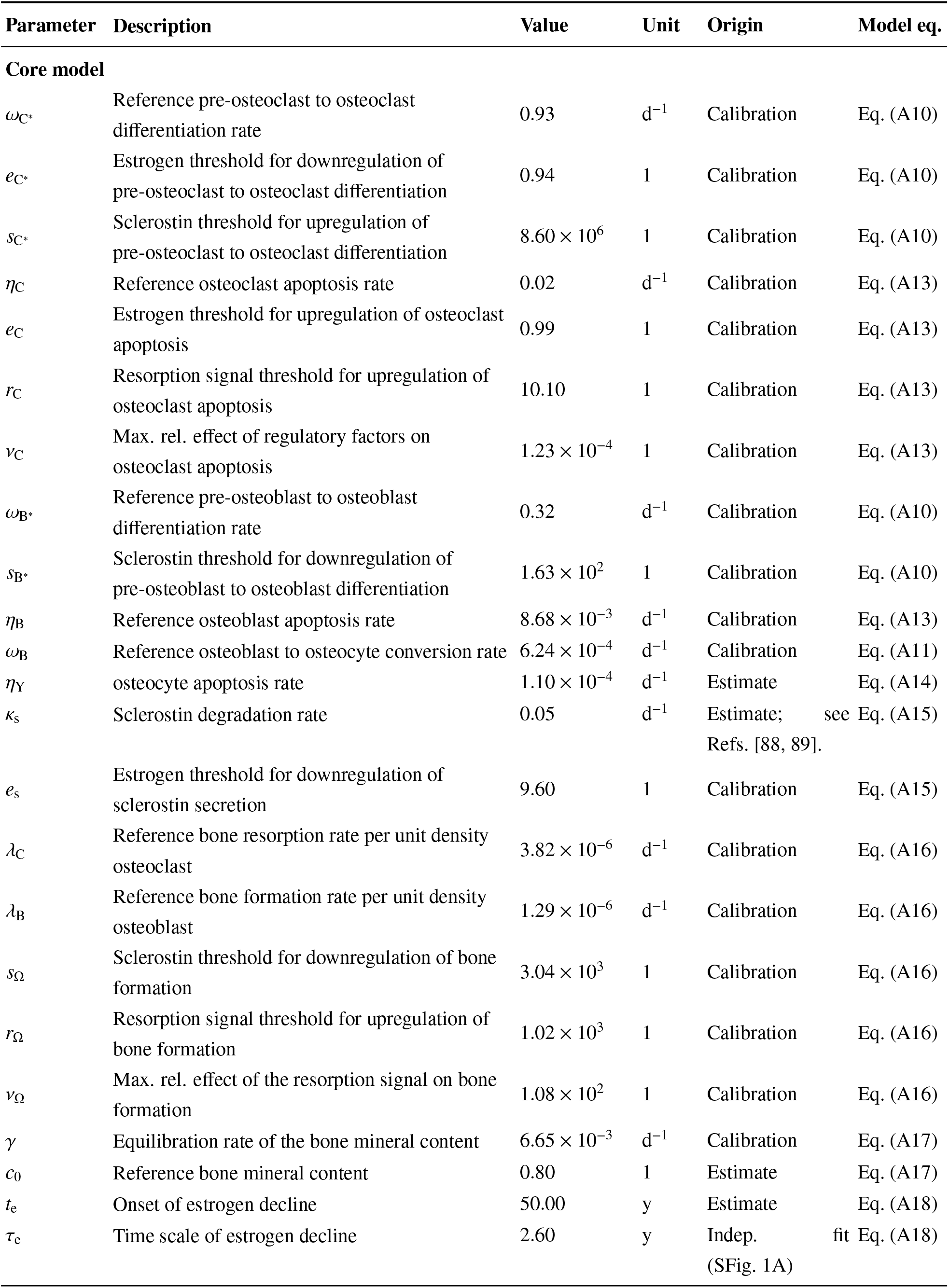
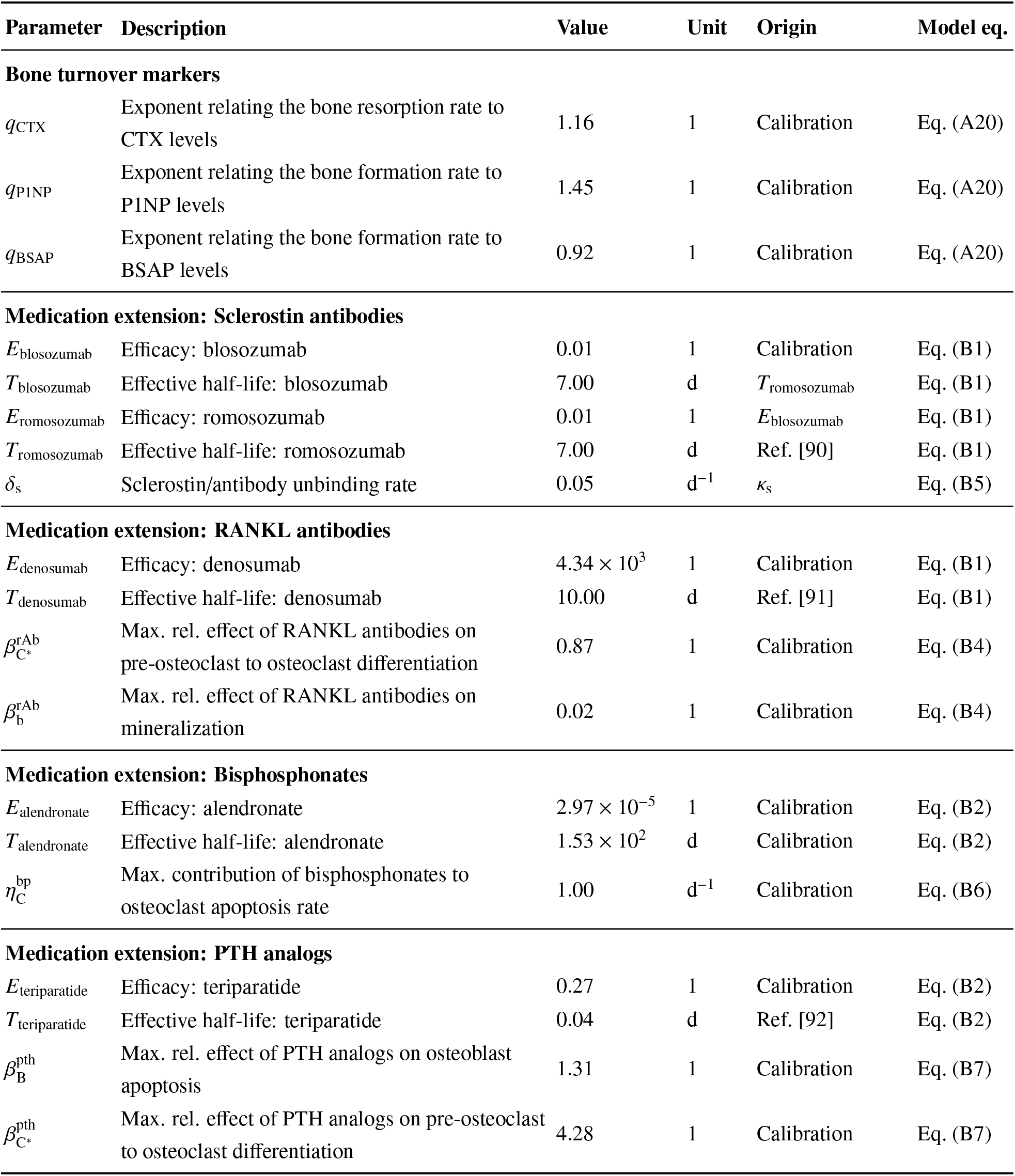
Full list of parameters of the core model and the medication extensions.

| Parameter | Description | Value | Unit | Origin | Model eq. |
| --- | --- | --- | --- | --- | --- |
| **Core model** |  |  |  |  |  |
| $\omega_{C^*}$ | Reference pre-osteoclast to osteoclast differentiation rate | 0.93 | d<sup>-1</sup> | Calibration | Eq. (A10) |
| $e_{C^*}$ | Estrogen threshold for downregulation of pre-osteoclast to osteoclast differentiation | 0.94 | 1 | Calibration | Eq. (A10) |
| $s_{C^*}$ | Sclerostin threshold for upregulation of pre-osteoclast to osteoclast differentiation | $8.60 \times 10^6$ | 1 | Calibration | Eq. (A10) |
| $\eta_C$ | Reference osteoclast apoptosis rate | 0.02 | d<sup>-1</sup> | Calibration | Eq. (A13) |
| $e_C$ | Estrogen threshold for upregulation of osteoclast apoptosis | 0.99 | 1 | Calibration | Eq. (A13) |
| $r_C$ | Resorption signal threshold for upregulation of osteoclast apoptosis | 10.10 | 1 | Calibration | Eq. (A13) |
| $\nu_C$ | Max. rel. effect of regulatory factors on osteoclast apoptosis | $1.23 \times 10^{-4}$ | 1 | Calibration | Eq. (A13) |
| $\omega_{B^*}$ | Reference pre-osteoblast to osteoblast differentiation rate | 0.32 | d<sup>-1</sup> | Calibration | Eq. (A10) |
| $s_{B^*}$ | Sclerostin threshold for downregulation of pre-osteoblast to osteoblast differentiation | $1.63 \times 10^2$ | 1 | Calibration | Eq. (A10) |
| $\eta_B$ | Reference osteoblast apoptosis rate | $8.68 \times 10^{-3}$ | d<sup>-1</sup> | Calibration | Eq. (A13) |
| $\omega_B$ | Reference osteoblast to osteocyte conversion rate | $6.24 \times 10^{-4}$ | d<sup>-1</sup> | Calibration | Eq. (A11) |
| $\eta_Y$ | osteocyte apoptosis rate | $1.10 \times 10^{-4}$ | d<sup>-1</sup> | Estimate | Eq. (A14) |
| $\kappa_s$ | Sclerostin degradation rate | 0.05 | d<sup>-1</sup> | Estimate; see Refs. [88, 89]. | Eq. (A15) |
| $e_s$ | Estrogen threshold for downregulation of sclerostin secretion | 9.60 | 1 | Calibration | Eq. (A15) |
| $\lambda_C$ | Reference bone resorption rate per unit density osteoclast | $3.82 \times 10^{-6}$ | d<sup>-1</sup> | Calibration | Eq. (A16) |
| $\lambda_B$ | Reference bone formation rate per unit density osteoblast | $1.29 \times 10^{-6}$ | d<sup>-1</sup> | Calibration | Eq. (A16) |
| $s_\Omega$ | Sclerostin threshold for downregulation of bone formation | $3.04 \times 10^3$ | 1 | Calibration | Eq. (A16) |
| $r_\Omega$ | Resorption signal threshold for upregulation of bone formation | $1.02 \times 10^3$ | 1 | Calibration | Eq. (A16) |
| $\nu_\Omega$ | Max. rel. effect of the resorption signal on bone formation | $1.08 \times 10^2$ | 1 | Calibration | Eq. (A16) |
| $\gamma$ | Equilibration rate of the bone mineral content | $6.65 \times 10^{-3}$ | d<sup>-1</sup> | Calibration | Eq. (A17) |
| $c_0$ | Reference bone mineral content | 0.80 | 1 | Estimate | Eq. (A17) |
| $t_e$ | Onset of estrogen decline | 50.00 | y | Estimate | Eq. (A18) |
| $\tau_e$ | Time scale of estrogen decline | 2.60 | y | Indep. fit (SFig. 1A) | Eq. (A18) |
| **Bone turnover markers** |  |  |  |  |  |
| $q_{\text{CTX}}$ | Exponent relating the bone resorption rate to CTX levels | 1.16 | 1 | Calibration | Eq. (A20) |
| $q_{\text{PINP}}$ | Exponent relating the bone formation rate to PINP levels | 1.45 | 1 | Calibration | Eq. (A20) |
| $q_{\text{BSAP}}$ | Exponent relating the bone formation rate to BSAP levels | 0.92 | 1 | Calibration | Eq. (A20) |
| **Medication extension: Sclerostin antibodies** |  |  |  |  |  |
| $E_{\text{blosozumab}}$ | Efficacy: blosozumab | 0.01 | 1 | Calibration | Eq. (B1) |
| $T_{\text{blosozumab}}$ | Effective half-life: blosozumab | 7.00 | d | $T_{\text{romosozumab}}$ | Eq. (B1) |
| $E_{\text{romosozumab}}$ | Efficacy: romosozumab | 0.01 | 1 | $E_{\text{blosozumab}}$ | Eq. (B1) |
| $T_{\text{romosozumab}}$ | Effective half-life: romosozumab | 7.00 | d | Ref. [90] | Eq. (B1) |
| $\delta_s$ | Sclerostin/antibody unbinding rate | 0.05 | d<sup>-1</sup> | $\kappa_s$ | Eq. (B5) |
| **Medication extension: RANKL antibodies** |  |  |  |  |  |
| $E_{\text{denosumab}}$ | Efficacy: denosumab | $4.34 \times 10^3$ | 1 | Calibration | Eq. (B1) |
| $T_{\text{denosumab}}$ | Effective half-life: denosumab | 10.00 | d | Ref. [91] | Eq. (B1) |
| $\beta_{C^*}^{\text{rAb}}$ | Max. rel. effect of RANKL antibodies on pre-osteoclast to osteoclast differentiation | 0.87 | 1 | Calibration | Eq. (B4) |
| $\beta_b^{\text{rAb}}$ | Max. rel. effect of RANKL antibodies on mineralization | 0.02 | 1 | Calibration | Eq. (B4) |
| **Medication extension: Bisphosphonates** |  |  |  |  |  |
| $E_{\text{alendronate}}$ | Efficacy: alendronate | $2.97 \times 10^{-5}$ | 1 | Calibration | Eq. (B2) |
| $T_{\text{alendronate}}$ | Effective half-life: alendronate | $1.53 \times 10^2$ | d | Calibration | Eq. (B2) |
| $\eta_C^{\text{bp}}$ | Max. contribution of bisphosphonates to osteoclast apoptosis rate | 1.00 | d<sup>-1</sup> | Calibration | Eq. (B6) |
| **Medication extension: PTH analogs** |  |  |  |  |  |
| $E_{\text{teriparatide}}$ | Efficacy: teriparatide | 0.27 | 1 | Calibration | Eq. (B2) |
| $T_{\text{teriparatide}}$ | Effective half-life: teriparatide | 0.04 | d | Ref. [92] | Eq. (B2) |
| $\beta_B^{\text{pth}}$ | Max. rel. effect of PTH analogs on osteoblast apoptosis | 1.31 | 1 | Calibration | Eq. (B7) |
| $\beta_{C^*}^{\text{pth}}$ | Max. rel. effect of PTH analogs on pre-osteoclast to osteoclast differentiation | 4.28 | 1 | Calibration | Eq. (B7) |

### Goodness of fit measures

To assess the goodness of the parameter fit and model predictions, we considered complementary goodness measures. The mean absolute percentage error (MAPE) between clinical and simulated results is defined by

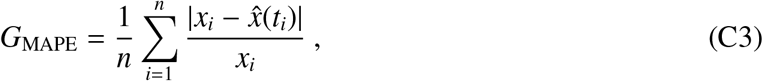

where the sum runs over all clinically recorded time points for the respective quantity (baseline changes of BMD and BTM levels), *x_i_* denotes the clinical data point and *t_i_* the time it was taken, 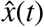 the model result, which is a continuous function of time.

As a complementary measure, we introduce a ‘windowed minimal absolute percentage error’ (WMAPE), which indicates the mean minimal distance between model results and the data within a time window around the data point that reflects the average time spacing between data points. Formally, the WMAPE is given by

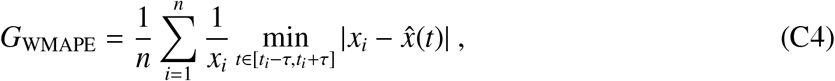

Here we choose the time window *τ* as half the median distance between data points, *τ* = median_*i*_(*t_i_* – *t*_*i*–1_)/2.

---

**Supplementary Figure 1.**
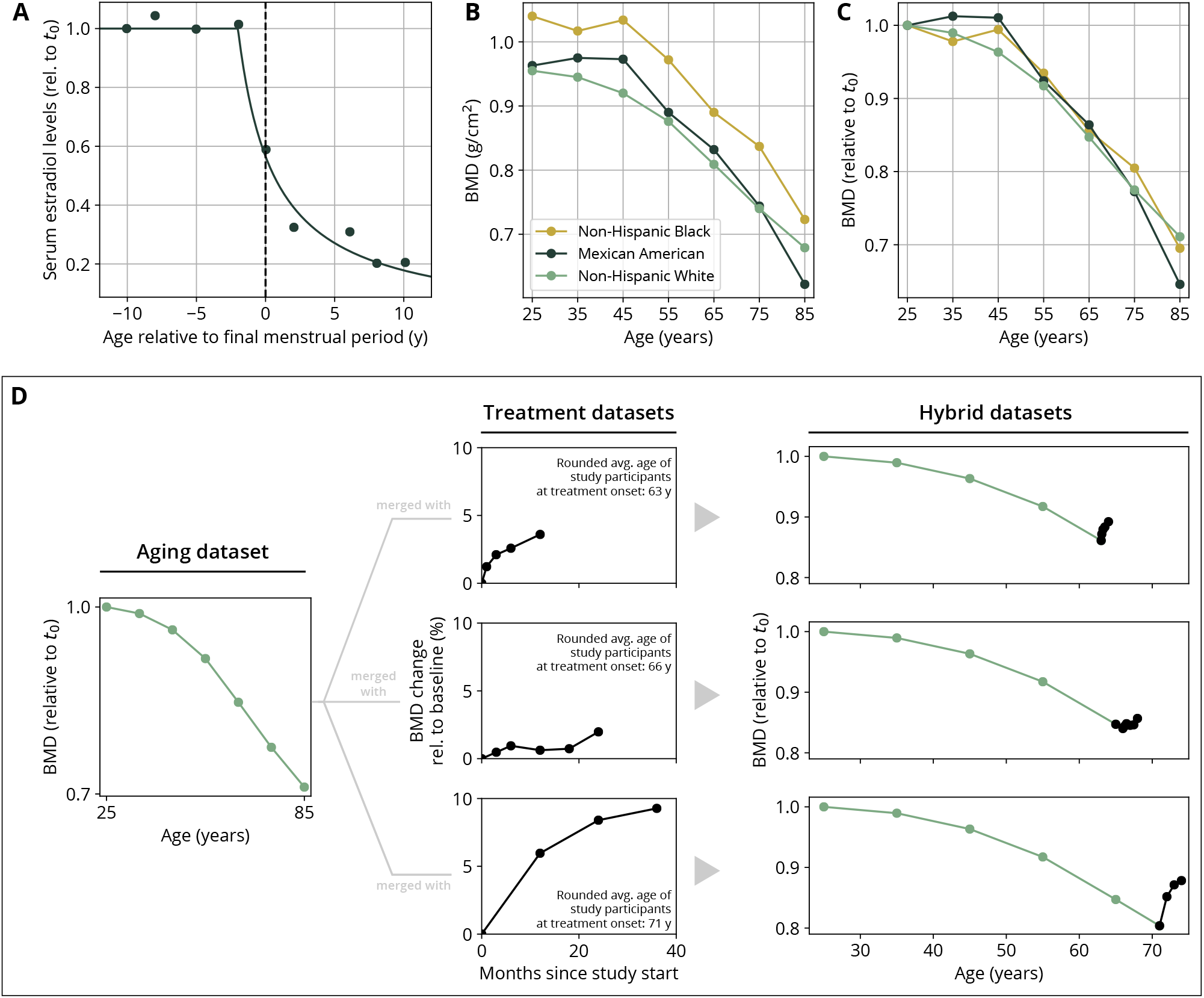
Parametrization of the aging behavior and creation of hybrid aging/treatment datasets for model calibration and validation. (A) Age dependence of estradiol serum levels. Clinical data (dots) modified from Sowers *et al*., 2008 [52]. The curve shows a fit of the function given by Eq. (A18) to determine the parameter *τ*_e_ (Table 4). (B) BMD age dependence for different ethnic groups as indicated. Data modified from Looker *et al*., 1998 [55]; reported age bin averages have been used to represent the center of the age bin. (C) BMD age dependence shown in panel B, where all curves have been normalized to their earliest value (*t*_0_ = 25y). (D) Schematic of how hybrid aging/treatment datasets were generated by merging the same aging dataset with different treatment datasets; for details, see Materials and Methods.

**Supplementary Figure 2.**
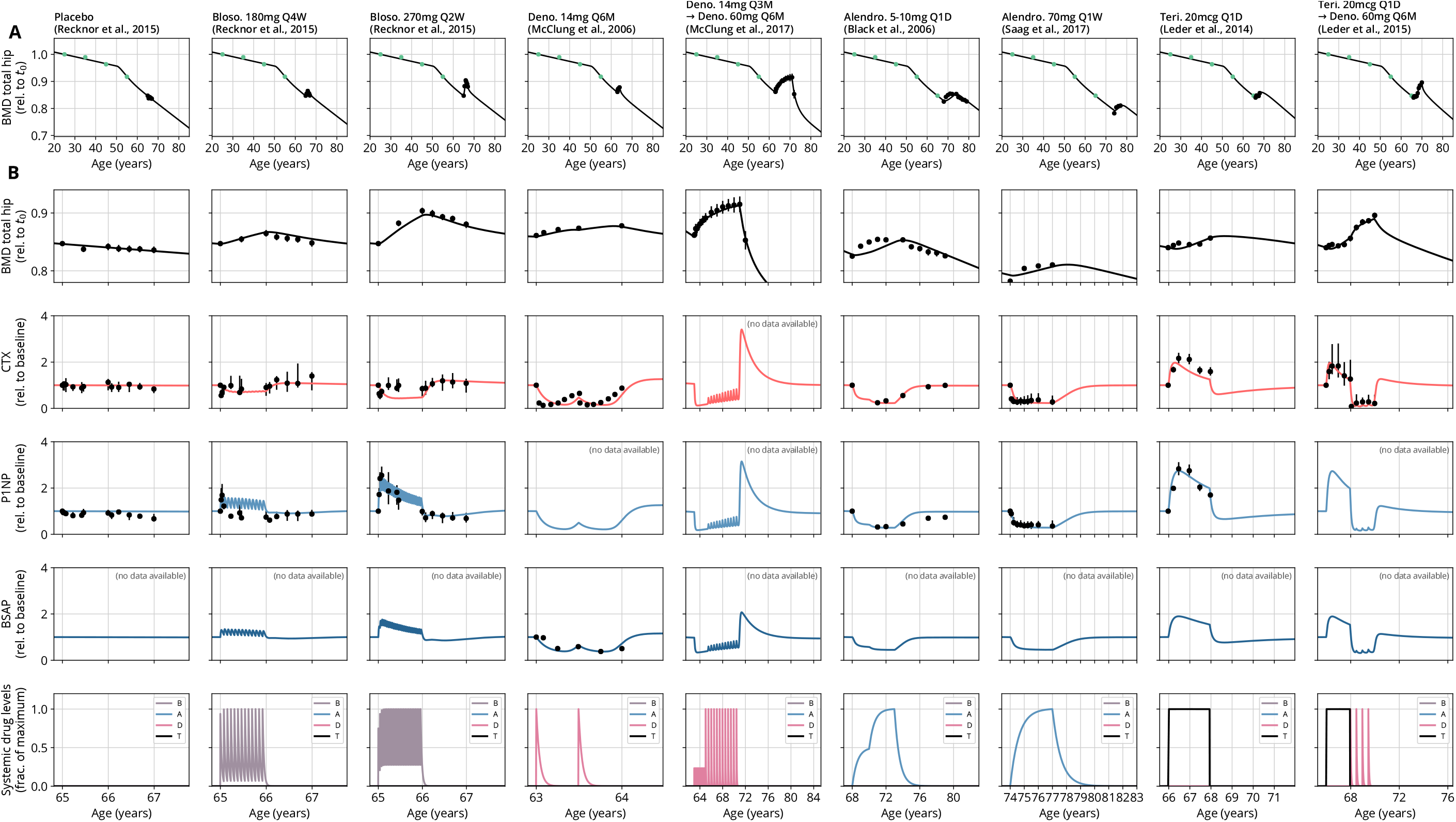
Calibration datasets comparing model predictions and clinical data from various studies. All conventions identical to main text Fig. 2. Drug administrations are provided in the bottom row. See Table 1 for a list of data sources and Table 2 for goodness-of-fit measures. Abbreviations (dosing): mg—milligrams; mcg—micrograms; Q*x*M—dose administered every *x* months; Q*x*W—every *x* weeks; Q*x*D—every *x* days. Abbreviations in bottom plot legend: B—blosozumab, A—alendronate, D—denosumab, T—teriparatide.

**Supplementary Figure 3.**
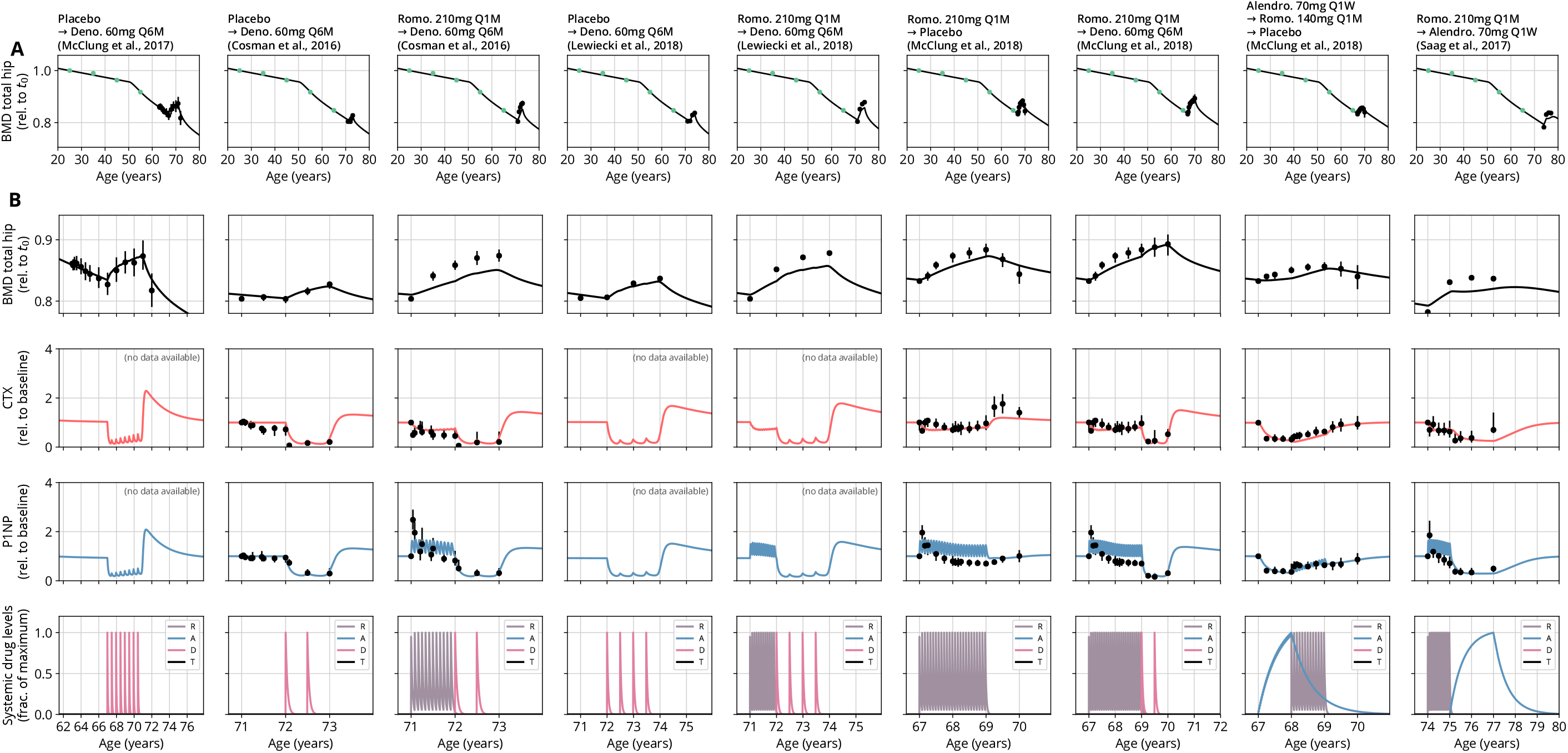
Validation datasets comparing model predictions and clinical data from various studies, continued from main text Fig. 2, all conventions identical. See Table 1 for a list of data sources and Table 2 for goodness of fit measures. Abbreviations (dosing): mg—milligrams; mcg—micrograms; Q*x*M—dose administered every *x* months; Q*x*W—every *x* weeks; Q*x*D—every *x* days. Abbreviations in bottom plot legend: R—romosozumab, A—alendronate, D—denosumab, T—teriparatide.

## Footnotes

1 Numerically, the quasi-continuous scheme has the advantage that model simulations do not have to resort to extremely small integration time steps to capture the details of short-term drug degradation, which considerably improves runtime.

